# Single-cell impedance cytometry of anticancer drug-treated tumor cells exhibiting mitotic arrest state to apoptosis using low-cost silver-PDMS microelectrodes†

**DOI:** 10.1101/2023.10.04.560818

**Authors:** Xinlong Yang, Ziheng Liang, Yuan Luo, Xueyuan Yuan, Yao Cai, Duli Yu, Xiaoxing Xing

## Abstract

Chemotherapeutic drugs such as paclitaxel and vinblastine interact with the microtubules, and thus induce complex cell states of mitosis arrest at the G2/M phase followed by apoptosis dependent on drug exposure time and concentration. Microfluidic impedance cytometry (MIC) as a label-free and high-throughput technology for single-cell analysis, has been applied for viability assay of cancer cells post drug exposure at fixed time and dosage, yet verification of this technique for varied tumor cell states after anticancer drug treatment remains vacant. Here we present a novel MIC device and for the first time perform impedance cytometry on carcinoma cells exhibiting progressive states of G2/M arrest followed by apoptosis related to drug concentration and exposure time, after treatments by paclitaxel and vinblastine, respectively. Our results from impedance cytometry reveal increased amplitude and negative phase shift at low frequency, as well as higher opacity for the Hela cells under G2/M mitotic arrest compared to the untreated cells. The cells under apoptosis, on the other hand, exhibit opposite changes in these electrical parameters. Therefore, the impedance features differentiate the Hela cells under progressive states post anticancer drug treatment. We also demonstrate that vinblastine poses a more potent drug effect than paclitaxel especially at low concentrations. Our device is fabricated with a unique sacrificial layer-free soft lithography process as compared to the existing MIC device, which gives rise to readily aligned parallel microelectrodes made of silver-PDMS embedded in PDMS channel sidewalls with one molding step. Our results uncover the potential of the MIC device, with a fairly simple and low-cost fabrication process, for cellular state screening in anticancer drug therapy.

## Introduction

Cancer has become a global burden with rapidly growing incidence and mortality.^1^ Chemotherapy is one of the main options for cancer treatment other than surgical resection and radiotherapy, where cytotoxic agents are given to kill the tumor cells or inhibit their proliferation.^2^ Therefore, investigation of the cellular response to an anticancer drug is of great significance not only for drug discovery regarding evaluating the effectiveness of a new drug, but also for monitoring the therapy response to facilitate personalized medicine. Conventional analytical approaches assess the viability of a cell population using microscopic inspection or colorimetric assays, such as trypan blue inclusion of cell membrane integrity, the terminal transferase-mediated dUTP nick end-labelling (TUNEL) assay detecting DNA fragmentation, or MTT assay for cell metabolic activity. Flow cytometry as a more advanced technique enables differentiation between early and late-stage apoptosis at single-cell resolution via fluorescence labelling of Annexin V and cell nuclear.^3^ However, it involves costly dyes as well as cumbersome labelling procedures.

Microfluidic platforms screening the electrophysiology status of anticancer drug-treated cells have been increasingly investigated in the last decade due to their label-free and non-invasive nature. Dielectrophoresis (DEP) is a commonly used electrokinetic method that measures the averaged cell dielectric properties according to the response of a cell population to a non-uniform electric field at varying frequencies. It has been demonstrated to be able to detect apoptosis induced by anticancer drug earlier and more rapidly than the conventional staining-based process.^3,4^ In addition to “bulk assays” on cell populations, Li et al. reported a single-cell level study using DEP for the chemotherapeutic resistance heterogeneity of leukemic blasts.^5^ Similarly, electro-rotation (ER) that extracts the dielectric properties of individual cells by measuring their frequency-dependent rotation speed has also been exploited in characterizing the multi-drug resistance of cancer cells.^6^ One challenging issue for these two electrokinetic methods is the precise positioning of individual cells in the analysis region near the activation microelectrode, which limits their application in high throughput analysis of single cells.^5-7^ Huang et al. improved the cell loading stability by applying DEP manipulation for ER chip.^8^

Microfluidic impedance cytometry (MIC) enables label-free single cells analysis with comparably high throughput to flow cytometry, i.e. > 10,000 cells in a few minutes.^9, 10^ It directly measures the electric current change as a cell passes through a pair of microelectrodes under electrical activation at single or multiple frequencies. MIC has been exploited for single-cell analysis of plant cells,^11^ parasites,^12^ fungi,^13^ stem cells,^14^ cancer cells,^7^ blood cells,^15,16^ etc. It maps cell biophysical properties, such as cell size,^17^ membrane capacitance and cytoplasm conductivity,^10^ or even mechanical deformability,^18^ to cell subpopulations with different types or cellular states.^13,18,19,20^ Recent review papers give very good summaries regarding works that map cellular/subcellular impedance features to phenotyping applications.^21,22^

Researchers in recent years start the investigations of MIC devices for the characterization of anticancer drug-treated cancer cells using MIC devices. Svendsen et al. exploited the MIC device with a three-electrode coplanar configuration to distinguish between the paclitaxel-treated and non-treated Hela cells, yet at a fairly low throughput of ∼3 cells/s.^23^ Ninno et al. applied a similar configuration for high throughput (>200 cells/s) discrimination of viable, apoptotic and necrotic human lymphoma U937 cells after being subject to etoposide treatment or heat shock.^24^ Gong et al. recently reported monitoring of paclitaxel-treated cancer spheroids for viability assessment.^25^ Honrado et al. measured the gemcitabine-induced apoptosis of pancreatic tumor cells.^26^ These existing works focus more on the viability analysis of tumor cells post anticancer drug treatment. However, cellular response to a given chemotherapeutic drug can be complex. For instance, paclitaxel (taxol) and vinblastine (VBL), commonly used broadest-spectrum antineoplastic agents, have been shown to induce cell mitosis arrest at G2/M phase cycles followed by cell apoptosis, as shown in Fig. 1(a), and the cell states progression exhibits dependence on the concentration and duration of drug exposure.^27-29^ MIC devices potentially uncover changed properties of tumor cells regarding size, morphology as well as electrophysiology properties as a response to the exposure time and concentration of anticancer drug treatment other than cell death that has been investigated by existing devices.

**Fig. 1.**
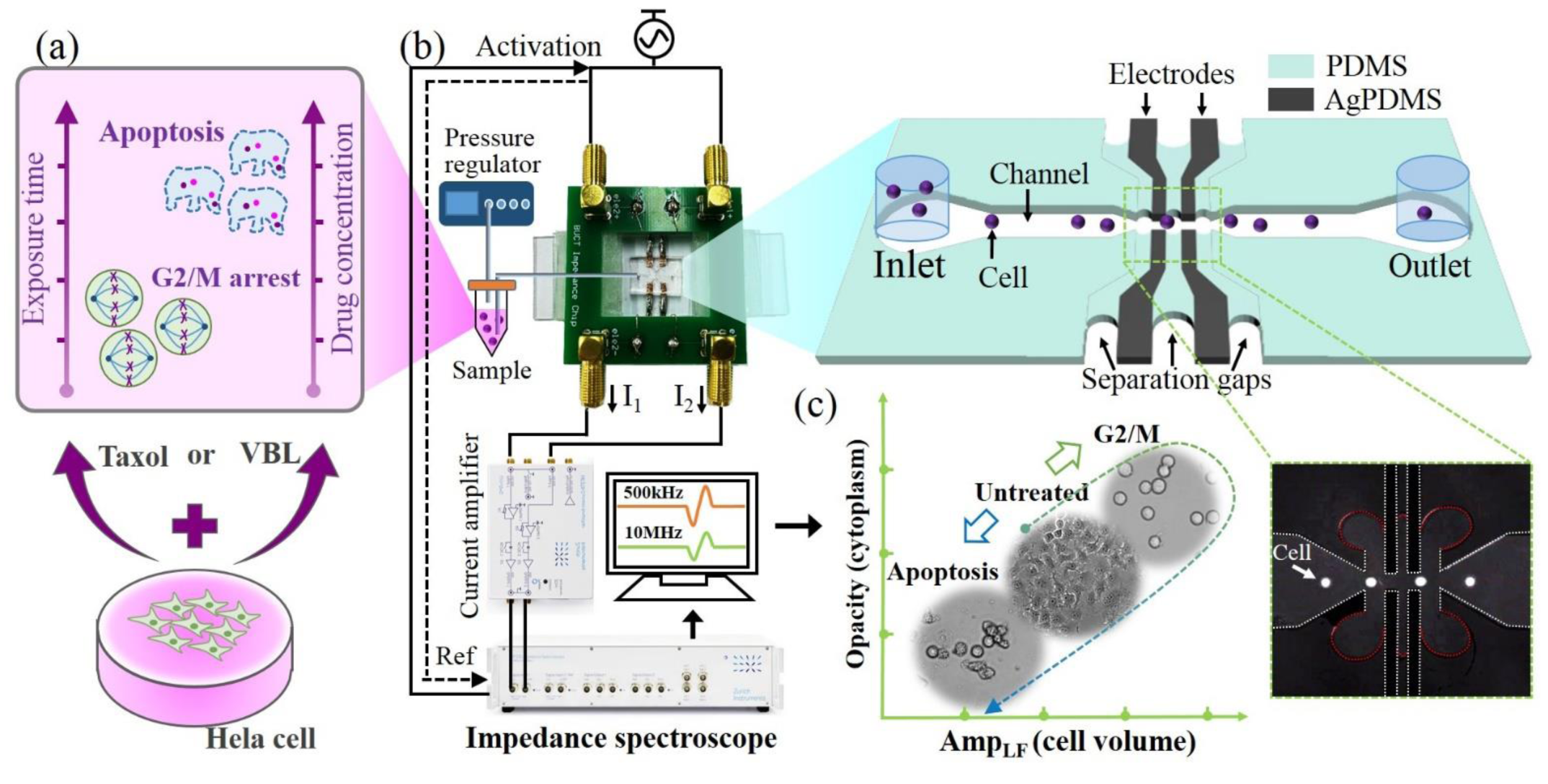
(a) Schematic illustration of Hela cells treated with taxol or VBL that exhibit drug concentration- and exposure time-dependent cell states of G2/M arrest and apoptosis. (b) Overview of the system setup and 3D schematic of the device design. Inset image in the bottom right corner: superimposed image of cell trajectory passing through the measurement region. White dashed lines: channel and electrodes boundaries. Red dashed lines: interfaces of capillary-filled PDMS. (c) Illustration of the microscopic images exhibiting cell states differentiated by the measured impedance data (i.e. opacity and low frequency amplitude (Amp^LF^) that reveal the cell volume and cytoplasm properties). The images are cells in culture dish under untreated, G2/M arrest and apoptosis states, respectively.

On the other hand, electrode configurations play a vital role in a MIC device, as have been reviewed comprehensively in the recent review papers.^30,31^ Coplanar metal electrodes patterned^11,15,16,20^ on the channel floor or parallel electrodes aligned face-to-face on the channel top and bottom^12,14,17,19^ are the most popular designs. Yet, these electrodes require standard cleanroom processes such as metal evaporation and lift-off, as well as aligned and leaky-free bonding, which gain the cost and complexity of fabrication. Alternatively, researchers report devices with simpler and more cost-effective processes of fabrication such as those exploiting soft lithography^10,25,32,33^ or laser cutting.^34^ For instance, Chen et al. presented high throughput characterization of the cell electrical properties using a PDMS-casted constriction channel configured with crossing side channels filled with ionic buffers to serve as liquid electrodes.^33^ Recently, Gong et al. reported the use of Field’s metal solidified in PDMS trenches that formed coplanar electrodes at the channel bottom.^25^ Yet, liquid electrodes or low melting point alloys are always concerned with the contamination or fabrication accuracy. In addition to soft lithography-based approaches, Ni et al. recently exploited low-cost ITO-coated polymer electrodes made of laser cutting,^34^ yet it again involved electrode alignment during the package. Therefore, low-cost and easily fabricated devices with high accuracy are still needed.^30^

In this work, we present an easily-fabricated MIC device featuring 3D monolithic silver-PDMS (AgPDMS) microelectrodes for high throughput single-cell analysis of anticancer drug-treated Hela cells. As shown in Fig. 1(b), the device has one straight flow channel that becomes constricted at the center portion. In the constricted region, two pairs of parallel-facing AgPDMS electrodes are embedded within the PDMS sidewalls and configured in differential mode. The AgPDMS electrodes and PDMS sidewall blocks are physically separated by narrow gaps filled with PDMS at the end of the fabrication process, which serve for electrical insulation and maintain the integrity of the entire device structure. The unique process used here allows the casting of the AgPDMS and PDMS blocks in one step by a readily-used mold without sacrificial layer lithography, which is typically required in existing devices incorporating conducting PDMS electrodes.^35^

In contrast to prior works, we use the device to investigate the concentration- and duration-dependent cellular responses of post drug-treated cancer cells that exhibit mitosis arrest at the G2/M phase as well as apoptosis, and two types of anti-tubulin chemotherapeutic agents, taxol and VBL, are respectively applied. 10,000 ∼ 15,000 cells are measured with a throughput of ∼250 cells/s. Impedance readouts are acquired at dual frequencies of 500 kHz and 10 MHz simultaneously for assessing the cell electrophysiology properties.^9^ We demonstrate that the measured amplitude, phase and opacity (i.e. the ratio of the high to low frequency amplitude), as indicators of the cell volume and dielectric properties, map the cells into states from untreated condition to progression stages of G2/M arrest and apoptosis (Fig. 1(c)). The results show the potential of this MIC device as a low-cost and label-free analytical tool for high-throughput single-cell analysis in anticancer drug screening.

## Materials and methods

### Microfabrication

Existing microfluidic devices featuring microelectrodes made of conducting PDMS (e.g. AgPDMS) all required additional sacrificial layer lithography on SU8 mold for separate patterning of the AgPDMS electrode and the PDMS flow sidewalls.^35,36^ In contrast, the MIC device here enables a simpler casting process free of sacrificial layer lithography. Fig. 2(a) schematically illustrates the fabrication process in a 3D view. The first step was photolithography of the mold, where the cavities designating the electrode as well as flow sidewalls were patterned within a 25 µm thick layer of SU8-2015 photoresist (MicroChem Corp, MA, USA) on a silicon wafer (Step i). The next step was replica molding. The AgPDMS composite was prepared by mixing the micro-sized silver particles (Xingrongyuan, Beijing, China) and PDMS gel (pre-mixed by 10:1 v/v of pre-polymer and curing agent, Dow Corning, MI, USA) at 85:15 weight ratio and then thoroughly grinded using pastel in a mortar to a homogeneous paste. The SU8 master treated with Trichloro (1H,1H,2H,2H-tridecafluoro-n-octyl) silane (T162729, Aladdin, Shanghai, China) was then filled substantially with the AgPDMS paste into the cavities (Step ii). Any excessive AgPDMS out of the cavities was cleaned with a piece of paper. Note that the mold cavities designating active and passive regions were simultaneously filled with AgPDMS in this step (Fig. S1(a); Active region: Region E denoting the electrodes; Passive region: Region W and P denoting the flow sidewalls and supporting posts, respectively.). After curing the AgPDMS at 70 °C for 3 hours, the cured AgPDMS thin films cured in the passive regions were peeled off by tweezer (Step iii, and a photo taken during this step was shown in Fig.S1(b)). These emptied regions were then filled by pouring PDMS onto the mold, where both the flow sidewalls (i.e. the passive region) and the device cap were formed (Step iv). The AgPDMS-PDMS layers were subsequently peeled off from the mold (Step v), punched with ports for fluids and PDMS filling, and boned onto a glass side with oxygen plasma treatment. During plasma bonding, the four AgPDMS electrodes respectively had contact with the metal stripes made of cooper tape sticking on the glass slide (Fig. S1(c)). The narrow trenches separating electrodes/flow sidewalls were capillary filled with half-cured PDMS, as shown in Fig. S2. The half-cured PDMS was prepared by heating the freshly-prepared PDMS at 70°C for 5 minutes to have it more viscous. The half-cured PDMS was dipped into the ports through the device layer that was aligned to those trenches. The filled PDMS was then cured at 120°C for 1 minute (Step vi). The chip was finally mounted onto a custom-printed circuit board (PCB) with soldering wires connecting each metal stripe to the PCB pad.

**Fig. 2.**
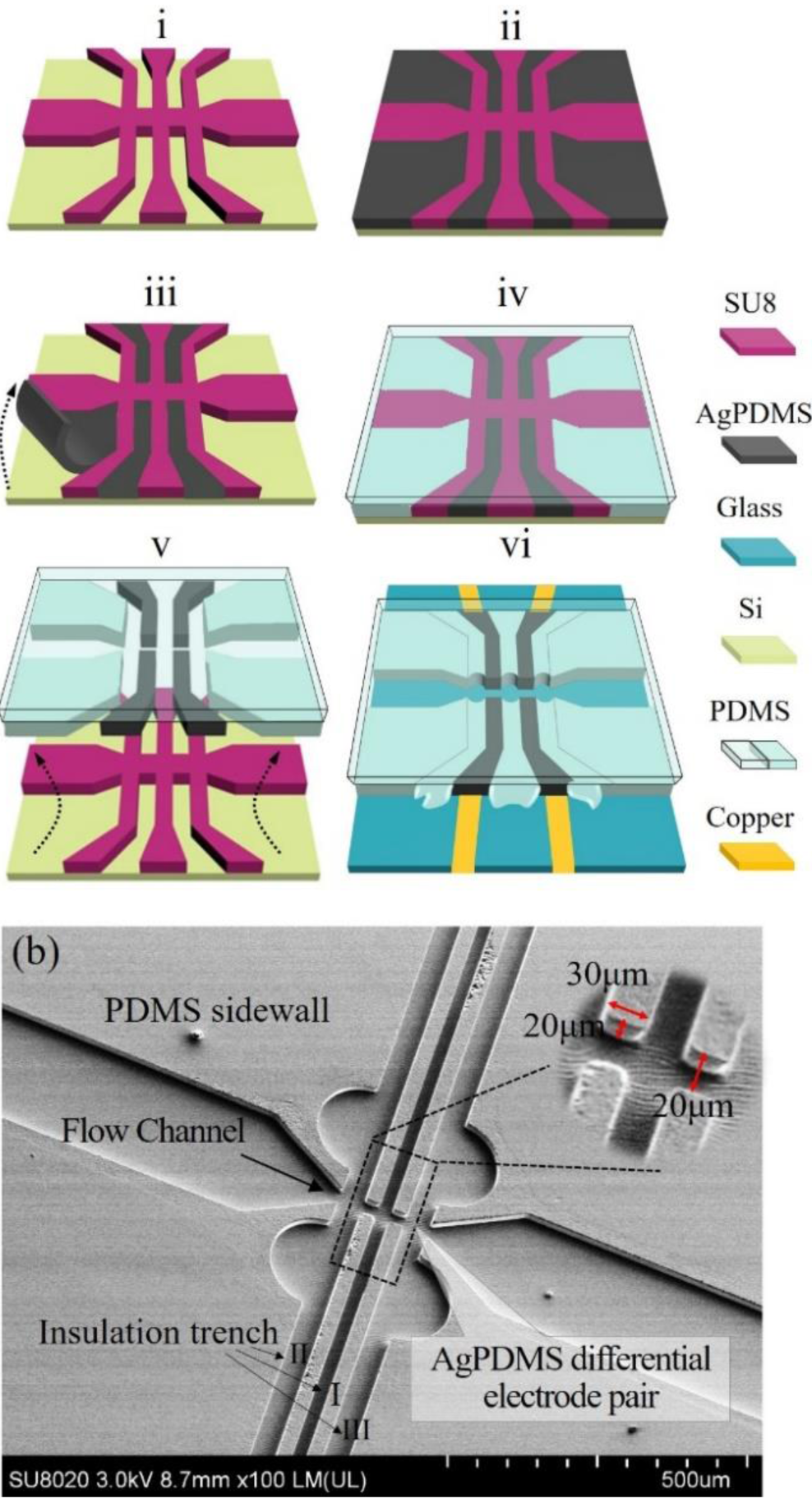
(a) Fabrication process flow illustrated by 3D schematics. (b) SEM image in oblique view for the fabricated device.

### Sample preparation

Hela cells in culture medium were cultured in a humidified incubator maintained at 37oC with 5% CO_2_. Before drug-treatment, the cells were seeded in a 90-mm tissue culture dish at 10^6^ cells per dish and pre-incubated for 24 h. Drug-containing culture mediums were prepared right before use by diluting the stock solution of taxol/VBL in fresh culture medium to designated concentrations. The cells were then treated with drug-containing culture mediums at 50-, 200-, 800-, and 3200-nM, and were incubated for 12-, 24-, 36- and 48-h. To avoid cell loss potentially caused by the cell detached from the dish after drug treatment, upon cell harvesting, the culture dish was flushed with the original culture medium in it to carry any suspended cells or any cells failing to attach firmly to the dish bottom (i.e., rounding cells at G2/M arrest state, apoptotic cells, or both). The cell-containing medium directly taken from the dish was then centrifuged together with the attached cells being harvested by standard trypsinization protocol, followed by being pooled together in fresh culture medium. Specifically, in the study of subpopulations of cells exhibiting rounding and flatting morphological features post 3200nM taxol treatment of 12h, the rounding cell contained in the original culture medium and the flatting cells attached at the dish bottom were collected separately without pooling. In addition to drug-treated cells, a control group with cells cultured in the absence of any drug was also harvested. Before the testing, the harvested cells were washed with PBS by centrifuge and finally re-suspended in PBS at the density of ∼2 × 10^6^ cells/ml.

### System setup for impedance cytometry

The system setup is shown in Fig. 1(b). The electrodes on one side of the channel were energized by sinusoidal voltage at dual-frequency of 500 kHz and 10 MHz and 1 V amplitude, from the function generator module of an impedance spectroscope (HF2IS, Zurich Instruments, Zurich, Switzerland). The electrodes on the other side of the channel measured the current in differential mode, followed by current to voltage conversion with 1 kΩ gain through a current amplifier (HF2TA, Zurich Instruments, Zurich, Switzerland). The signal was then demodulated by the impedance spectroscope, and the bandwidth for the low-pass filter of the lock-in amplifier was 1 kHz. The cell sample fed into the inlet was driven by regulated pressure of 10 mbar (OB1 MK3, Elveflow, Paris, France) with a high throughput of ∼250 cells/s. Successive passage of individual cells through the measurement region leads to the differential signal sequence being recorded under dual frequencies simultaneously, with a sampling rate of 57.6 kSa/s. The impedance readouts appear comparable with those measured using conventional coplanar electrodes regarding amplitude, transit time and throughput.^11,20^ An inverted fluorescence microscope (Ti2, Nikon, Tokyo, Japan) equipped with a CMOS camera (ORCA-Flash4.0, Hamamatsu, Tokyo, Japan) was used for image taking of the drug-treated cells in culture dish as well as monitoring the device during the experiment.

Reagents, processing algorithms and calibration of measured data, and operation approaches for flow cytometry can be found in the supplementary material.

## Results and discussion

### Device

Fig. 2(b) depicts the scanning electron microscopy (SEM) image of a replica molded device prior to bonding to the glass slide. The SEM image shows the main channel in oblique view being 20 µm deep and 200 µm wide, and it becomes constricted to be 20 µm wide at the centre measurement region, where the differential electrodes featuring 30 µm × 20 µm (width × height) cross-sectional area for each digit emerge embedded in the sidewall. The insulation trench “I” between two electrodes is 20 µm wide whereas the trench between the electrode and bulk flow sidewall (i.e. trench “II” or “III”) is 40 µm wide and expands to a semicircle shape near the joint to the main channel for the purpose of buffering the PDMS filled into these trenches. The differential sidewall electrodes appear to be readily aligned in the mask design. Such a design not only inherits the advantages of parallel-facing electrodes for having higher measurement sensitivity and less positional dependence on the measured particle, but it is also free of aligned bonding during fabrication. Moreover, it waives the need for separate lithography patterning of the flow channel and the electrode, which are typically required by MIC devices incorporating either metal electrodes^16,24,37^ or conducting PDMS-based electrodes in more recent works.^35^

In addition, we also measured the impedance spectra (Fig. S4) from one pair of the electrodes with the flow channel filled with deionized (DI) water and PBS respectively. The device with PBS exhibited a decreased impedance at relatively low frequency (∼<30kHz) due to the double layer capacitance. The decreasing trend is not significant in the figure due to the wide range of the y axis for incorporating the impedance curve of DI water. In higher frequency range, the device with PBS showed resistive plateau covering the working frequencies (i.e., 500 kHz and 10MHz) applied to the impedance analysis in this work.

### Impedance analysis of taxol-induced cell volume change in varied cell states

Taxol is a heavily used anticancer drug and it targets on cell tubulin and leads to the stabilization of the spindle microtubule. It has been well-documented that taxol induces progressive cell states of mitotic arrest at metaphase (traditionally also called G2/M arrest), followed by apoptotic cell death as a consequence of abnormal mitosis exit.^38-41^ Here, we first analyzed the cell electrical volume using the low frequency amplitude (i.e. ***Amp***_**500*k***_), being analogous to the coulter principle, as an indicator for the varied cellular states post taxol exposure respectively at concentrations of 50-, 200-, 800- and 3200-nM and durations of 12-, 24-, 36- and 48-h. Fig. 3(a) illustrates the time course of the histogram sets with respect to the applied drug concentrations. Each histogram displays the electrical volume distribution of 15,000 cell events, and its mean value with standard deviation is further extracted and presented in the bar plot in Fig. 3(b).

**Fig. 3.**
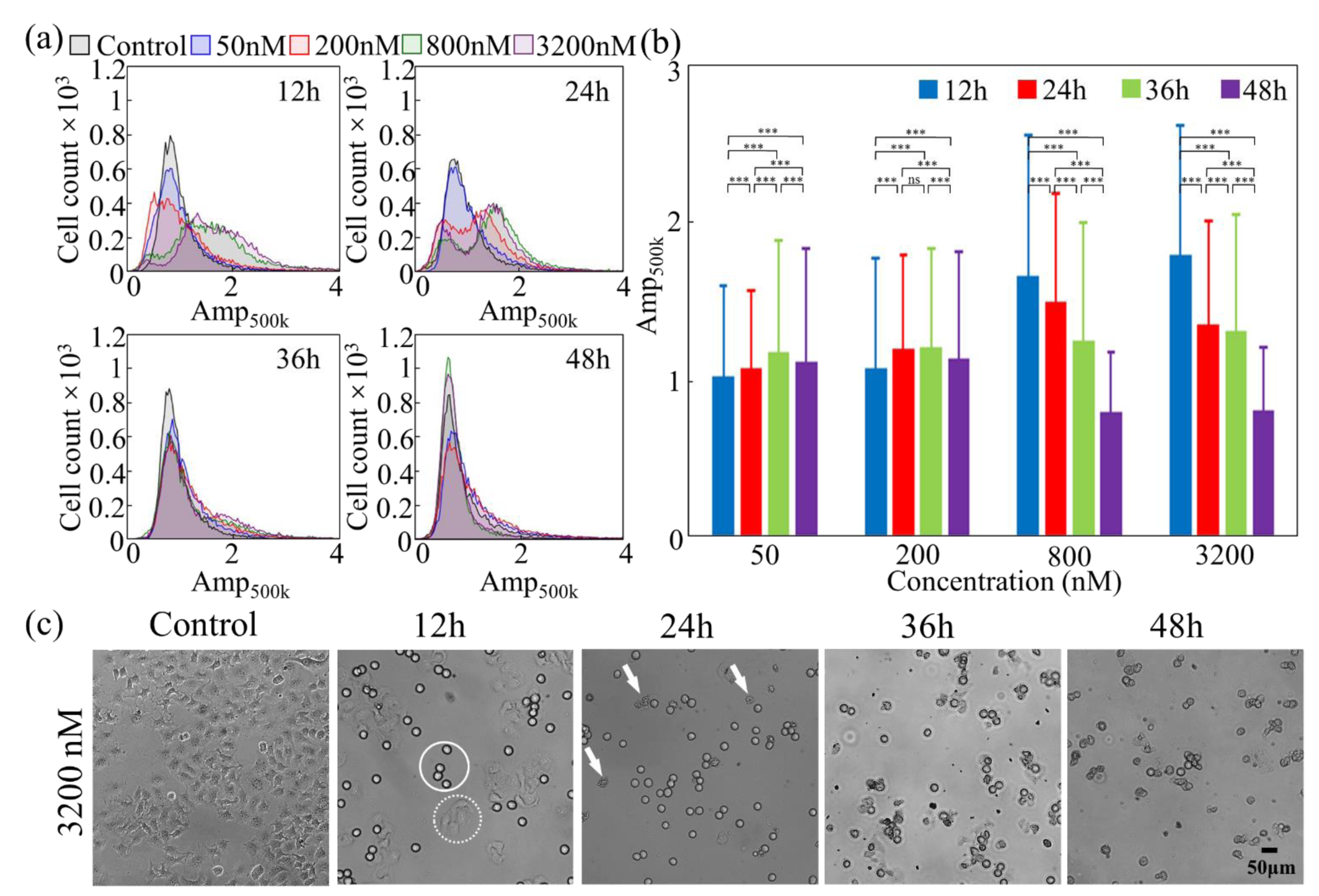
(a) Histograms of the low frequency amplitude (*AmP*_500*k*_) for cells post exposure to varying taxol concentrations of 0 (control), 50-, 200-, 800- and 3200-nM, respectively at the given exposure time durations of 12-, 24-, 36- and 48-h. Each histogram shows distribution of 15,000 events. (b) Mean ± s.d. of the *AmP*_500*k*_ extracted from each condition in (a) and plotted against the exposure time durations respectively under taxol concentrations of 50-, 200-, 800- and 3200-nM (n = 15,000). (c) Phase contrast images of the untreated cells and taxol-treated cells after the given exposure time durations (12-, 24-, 36- and 48-h) at 3200nM concentration. Outlined cells: representative cells exhibiting rounding (solid circle) and flatting (dashed circle) morphologies. Arrows: representative cells exhibiting blebbing membranes.

As revealed by Fig. 3(b), the electrical volume change post drug treatment exhibited an interesting pattern, where it expanded prior to the decrease. This is in contrast to the typical process of apoptotic volume decrease (AVD) for the anticancer-drug-treated cells, where they had volume shrinkage shortly after drug treatment (e.g. 30 minutes after staurosporine exposure).^42^ For instance, cells treated by 3200- and 800-nM taxol had peaked electric volume after a short exposure of 12 h, where the _**500*k***_ respectively exhibited a potent increase to 1.79 (3200nM) and 1.66 (800nM) with respect to the control group _**500*k***_ after calibration). As shown in Fig. 3(b), the _**500*k***_ then exhibited persistent decrease with prolonged exposure time intervals (i.e., 24-, 36- and 48-h). Such a trend was in agreement with the reported cell response to taxol,^40^ where the mitotic arrest led to cell volume increase at the beginning, followed by the decrease as a result of apoptosis after abnormal mitosis exit of these arrested cells.

We validate this by inspecting cell morphology features with phase contrast microscopic images, as demonstrated by the 3200nM taxol-treated cells in Fig. 3(c) (also see Fig. S5 in the supplementary materials). For 3200nM, cells after 12 h treatment exhibited a major population with an apparent rounding feature (outlined by a solid circle) as a signature of mitotic arrest at metaphase, which is also known as “mitotic rounding”^43^ with peaked volume in mitosis.^44^ The rest portion of the treated cells stayed flatting (outlined by a dashed circle) on the dish bottom. Starting from 24 h, as shown in Fig. 3(c), cell shrinkage occurred with membrane blebbing (pointed by arrows) as indicators of apoptosis, which gradually became severe with prolonged time length until 48 h. We also looked into cells exhibiting different morphological features separately (e.g. flatting and rounding) and validated using flow cytometry, as detailed later in the next section.

In comparison, cells after exposure to relatively lower taxol concentrations of 200- and 50-nM had rather moderate electrical volume increase, with the maximum ***Amp***_**500*k***_ being 1.20 (200nM) and 1.17 (50 nM) occurred at 36 h. These lowered ***Amp***_**500*k***_ peak values with delayed occurrence time when drug concentrations became lower (i.e. 200- and 50-nM) possibly related to the acceleration effect of taxol on cell cycle progression previously described.^45^ Treatment at higher taxol concentrations led to more rapid cell arrest at the metaphase during mitosis and, followed by more rapid apoptosis consequently. The microscopic images taken for these lower concentration conditions (i.e. 200- and 50-nM) also revealed much less effect of the drugs on cells, as shown in Fig. S5.

The volume change for mammalian cells experiencing different cell cycles in proliferation or undergoing apoptosis are also well-known morphological characteristics.^44,46,47^ During proliferation, cells in metaphase have significantly larger sizes than those at the G1/S phase. For instance, Rimington et al. reported ∼40% volume increase of Hela cells (ovarian cancer cell line) in the late G2-phase passing into M-phase than those in G1, and the cell volume in metaphase is twice that in early G1.^46^ Zlotkiewicz et al. reported that the volume increase of Hela cells is accompanied with mitosis until metaphase and the increase is independent of cell shape (i.e. being adherent or prorounded).^44^ Sugano et al. measured 24% mean diameter increase of G2/M cell cycle arrest for colon cancer cells (HCT116) as compared to those in the control group with regular cell cycle distribution.^48^ Kim et al. reported significant cell volume growth of human breast carcinoma cells (MDA-MB-231) from an average diameter of 10 µm in the G1/S phase to 20 µm in the G2/M phase.^49^ On the other hand, apoptosis, known as programmed cell death, causes persistent cell shrinkage and subsequent membrane blebbing to tiny apoptotic bodies.^20,42,47^

### Impedance analysis of subpopulations of G2/M arrested cells exhibiting different morphological features

We further investigated the volume change separately for the two apparent morphological features, i.e., rounding and flatting, exhibited in cell groups post 12 h treatment of 3200nM taxol (Fig. 3(c)), where they mostly underwent mitotic arrest process. The electrical volume of the rounding cells, the flatting cells, and the untreated cells were respectively measured using impedance cytometry. Validation by flow cytometry was also run in parallel. Scatter plot (Fig. 4(a)) of the ***Amp***_**500*k***_ against ***phase***_**500*k***_ displayed successive electrical volume increases of the flatting and rounding cells. Either two cell groups of the cell populations denoting the control, flatting and rounding cells exhibited statistically highly significant differences (P ≤ 0.001) for both the ***Amp***_**500*k***_ and ***phase***_**500*k***_ (Table S2). The histograms (Fig. 4(b)) of the ***Amp***_**500*k***_ distribution labelled with the mean value further revealed 59% and 94% increase of the mean electrical volume respectively for the flatting and rounding cells, which corresponded to 17% (flatting) and 25% (rounding) diameter increase by taking the cube root of the electrical volume. As validated by flow cytometry, the FSC-H data denoting cell diameter (Fig. 4(c)) also consistently exhibited an increasing trend for the flatting and rounding cells with diameter increments respectively at 14% and 18%. The significantly increased volumes suggested a mitotic arrest state not only for the rounding cells, but also for the flatting cells.

**Fig. 4.**
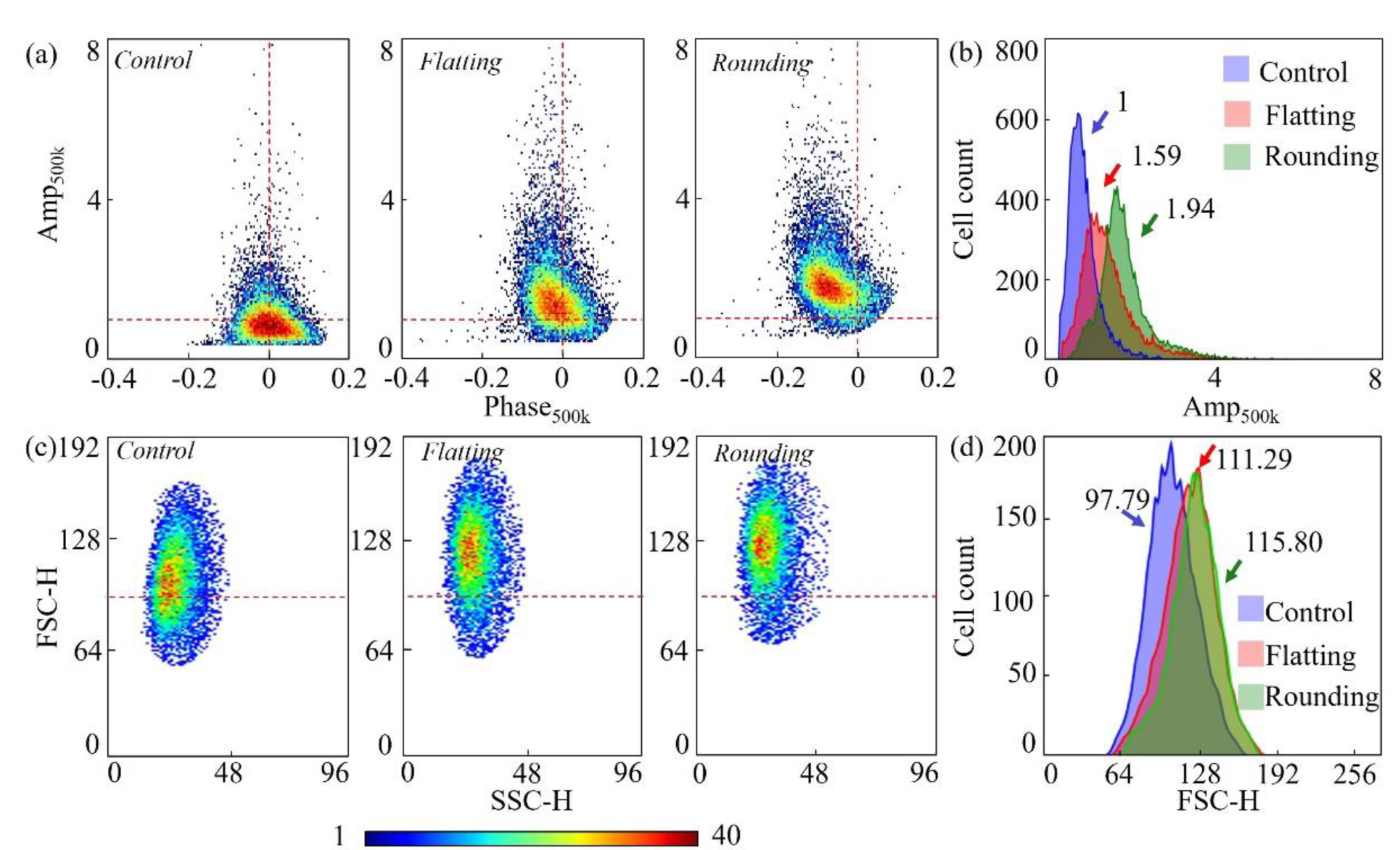
Analysis by (a-b) microfluidic impedance cytometry and (c-d) flow cytometry for the cell subpopulations respectively exhibiting flatting and rounding morphological features after taxol treatment at 3200 nM and 12 h as well as for the untreated cells (control). Impedance cytometry analysis displays (a) the scatter plots of low frequency amplitude (*Amp*_500*k*_) against phase (*Phase*_500*k*_) and (b) the histograms of *Amp*_500*k*_ for the control, flatting and rounding cells. Flow cytometry analysis presents (c) the scatter plots of the FSC‐H versus SSC‐H and the histograms of FSC‐H value for the control, flatting and rounding cells. Each scatter/histogram plot contains 10,000 cell events. The color bar designates the cell density at each data point.

Previous investigations^44,50,51^ had verified that cells experienced sharp volume growth from mitosis entry, also known as “mitotic swelling”, and reached a maximum during metaphase. Therefore, the even greater size of the rounding cells than the flatting ones indicated a more progressive mitosis stage in or approaching the metaphase of this cell population and, meanwhile, the mitotic rounding was also a morphological fingerprint for cells under these mitotic phases. The flatting cells, in contrast, designated those undergoing mitosis yet at a relatively early phase (e.g. prophase) without rounding up.

In addition to the volume increase, the scattered clusters of the flatting and rounding cells also exhibited successive negative phase shifts with respect to that of the untreated cells as designated by ***phase***_**500*k***_in Fig. 4(a). The phase signature at 500 kHz frequency arose from the capacitive behaviour of the cell membrane.^52^ The negative phase shift here was most probably due to the membrane capacitance increase, induced by the enlarged membrane area of these volume-expanding cells under mitotic arrest. Such a negative shift trend of phase also aligned with the modelled results.^9,20,53^ For an even longer treatment time under 3200nM until 48 h, we measured vanishing phase shift with apoptotic volume decrease, as illustrated in Fig. S8 of the supplementary material. Such vanishing phase shift corresponded to the reduction of the membrane capacitance possibly induced by the cell membrane perforation as well as shrunken membrane surface area during apoptosis.^23,54^

### Opacity of the taxol-treated cells under different states

At the high frequency of 10 MHz, the activation signal penetrated the cell membrane due to short-circuit of the capacitive membrane and thus the impedance measurement screened the electrophysiology feature of the cell interior (i.e. the cytoplasm conductivity).^52^ Here, we exploited opacity, denoted by ***Amp***_**10*m***_***Amp***_**500*k***_, for analysis of cell interior property during the cell state progression post taxol treatment.

Fig. 5 illustrated the scatter plot of opacity versus the ***Amp***_**500*k***_ for cells post 3200 nM taxol exposure for 12 h, 24 h and 36 h. In contrast to 12 h, where the majority of the cell population exhibited electrical volume growth (***Amp***_**500*k***_ > **1**), the cell events at 24 h evolved into two distinguishable subpopulations as could be divided by the normalized line of the electrical volume (***Amp***_**500*k***_= **1**). Such transition of distribution from 12 h to 24 h, with the appearance of the subpopulation featuring declined electrical volume (***Amp***_**500*k***_ < **1**), aligned with the volume decrease from 12 h to 24 h for 3200nM treated cells shown in Fig. 3(b), and it further verified the apoptotic cell state by the electrical volume shrinkage. The micrograph starting from 24 h (Fig. 3(c)) also displayed a noticeable amount of cells with membrane blebbing as a morphological signature of apoptosis other than those with mitotic rounding feature.

**Fig. 5.**
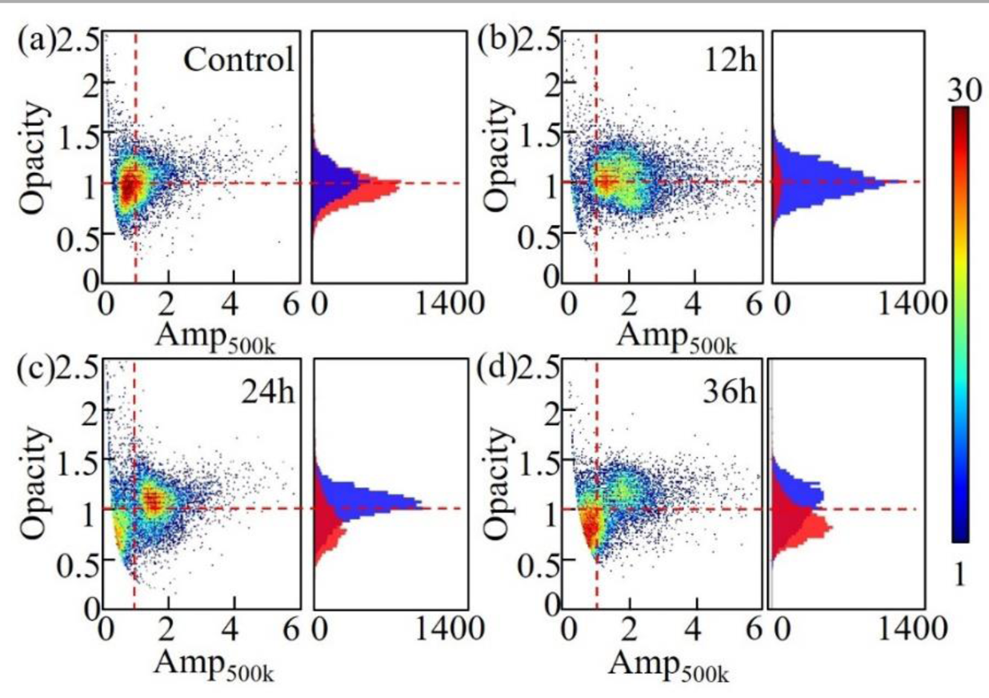
Scatter plots of opacity versus ***Amp***_**500*k***_ for (a) the untreated cells as control and (b-d) the cells treated by 3200 nM taxol respectively at (b) 12 h, (c) 24 h and (d) 36 h. The coloured histograms in each subfigure of (a)∼(d) show the opacity distribution respectively for cell subpopulations being classified by ***Amp***_**500*k***_ < **1** (blue) and ***Amp***_**500*k***_ < **1** (red). Each scatter plot contains 15,000 cell events. The color bar designates the cell density at each data point.

Coincident with the electrical volume distribution, the opacity here also exhibited a responsive pattern, as illustrated by the scatter plots and histograms of the volume-based subpopulations (Fig. 5). The opacity of the control group was normalized to distribute around 1 (**opacity** = **1**) after calibration. At 24 h, the subpopulation of ***Amp***_**500*k***_> **1** exhibited elevated opacity above the normalized line in the up-right region, as indicated by an up-shifted blue peak in the histogram, whereas those with ***Amp***_**500*k***_ < **1** had declined opacity below the normalized line in the down-left region, as shown by the down-shifted red peak. The cell population at 36 h exhibited similar subpopulation distribution, except that more cell events transited from the up-right region (***Amp***_**500*k***_ > **1, opacity** > **1**) to the down-left region (***Amp***_**500*k***_ < **1, opacity** < **1**). The increased opacity for the subpopulation in the up-right region was in accordance with the so-called “osmotic swelling” as proposed by the early researchers,^43,44,51^ where the cells undergoing mitotic volume growth had entry of pure water without accompany of increase in dry mass, and thus it led to diluted cytoplasm.^44,51^ Such dilution could lead to decreased cytoplasm conductivity that inversely resulted in more pronounced cell resistance at high frequency, as reflected here by the increased opacity. The “osmotic swelling” was a hypothesized mechanism that drove the mitotic rounding of the volume-expanded cells, which had been a debating point for long time.^44^ Our device here supports this hypothesis with the impedance characteristics, revealing the increased electrical volume accompanied by decreased cytoplasm conductivity with the coupled ***Amp***_**500*k***_ and opacity. On the contrary, the declined opacity of the other subpopulation in the down-left region aligned with cellular change during apoptosis. Cells under apoptosis exhibited perforated membrane and thus failed to maintain the contrast between the cytoplasm and the high conductivity suspension buffer,^23,54^ leading to lower resistance and thus lower opacity of the cells at high frequency.^17,54^ Statistically, the differences in the opacity between the two subpopulations of ***Amp***_**500*k***_ > **1** (blue peak) and ***Amp***_**500*k***_ < **1** (red peak) were significant for 12h treatment and highly significant for 24 and 36h treatments. The differences of opacity as well as ***Amp***_**500*k***_ were all highly significant between the two subpopulations in the up-right and down-left regions from 12h to 36h treatments (Table S3).

Flow cytometry has been used to validate the states of the taxol-treated cells measured by our MIC device. Overall, the PI histograms exhibited an accumulation of cell events at the G2/M phase for the drug-treated groups compared to the control, and the scatter plots of the Annexin V/PI stain revealed an elevated number of apoptotic cells. Cells treated with 3200nM taxol for 24h were shown as representative in Fig. S7(a-b) (the middle column). Moreover, the fractions of the two major subpopulations designating G2/M arrest and apoptosis states, as respectively shown in the up-right and the down-left regions in opacity/***Amp***_**500*k***_ scatter plots in Fig. 5, have been compared with those derived from flow cytometry. As shown in Fig. S7(c), both the MIC device and flow cytometry exhibited an increased percentage of G2/M arrested cells post 12h and 24h treatment but a drop after 36h, as well as a significant increase of cell apoptosis starting from 24h. The percentage values also appear comparable. Note that the cell fraction obtained from flow cytometry for apoptosis state included both the early and late apoptosis. Considering the possible progression of apoptosis during the delivery of cell samples, the percentage values have been further calibrated by subtracting the apoptosis fraction of the control group.

### Impedance analysis of the VBL-treated cells

In addition to taxol, we also performed impedance cytometry on Hela cells treated by another type of widely-used chemotherapeutic drug, the vinblastine (VBL), which was analogous to taxol for being an anti-tubulin agent and leading to the mitotic arrest of cells at G2/M phase followed by apoptotic cell death.^32^ Yet, VBL worked oppositely against taxol regarding the functioning mechanism on microtubule, where the former suppressed while the latter promoted the microtubule dynamics.^55,56^ For comparison, here we applied the VBL to Hela cells with the same concentrations and time lengths as we had done with taxol. The electrical volume change of Hela cells as being reflected by the low frequency amplitude ***Amp***_**500*k***_, as well as the opacity that coupled with the high frequency information, both revealed more potent effects of low concentration VBL regarding inducing cell mitotic arrest and apoptosis. Such a trend aligned with the previous publications that reported lower effective concentrations of VBL than taxol.^38,56^

As shown in Fig. 6(a), the electrical volume pattern of VBL-treated cells displayed a significant discrepant to that of the taxol-treated cells (Fig. 3(a)), especially at the lower drug concentrations. At 50 nM, for instance, the VBL-treated cells exhibited a volume increase much higher than the taxol-treated ones (i.e. 1.46 vs. 1.02 for ***Amp***_**500*k***_) after a short treatment time length of 12 h. Later at 24 h, the maximum volume of VBL-treated cells appeared, not only higher (1.51 vs. 1.06 for ***Amp***_**500*k***_) but also earlier (24 h vs. 36 h) than the peaked volume of the taxol-treated cells. Such a trend implied that more cells post 50nM VBL treatment, as compared to taxol treatment, were arrested at the G2/M stage of mitosis where the cells had apparent volume growth. As validation, as shown in Fig. 6(b), the phase-contrast images of cells after drug exposure (50nM, 12 h and 24 h) respectively for VBL and taxol also coincided well with such trend, where more cells under VBL treatment exhibited morphological features of mitotic arrest for being rounding (outlined in a solid circle) or pre-rounding (outlined in a dashed circle). Impedance cytometry for drug-treated cells at 200 nM also complied with such a trend.

**Fig. 6.**
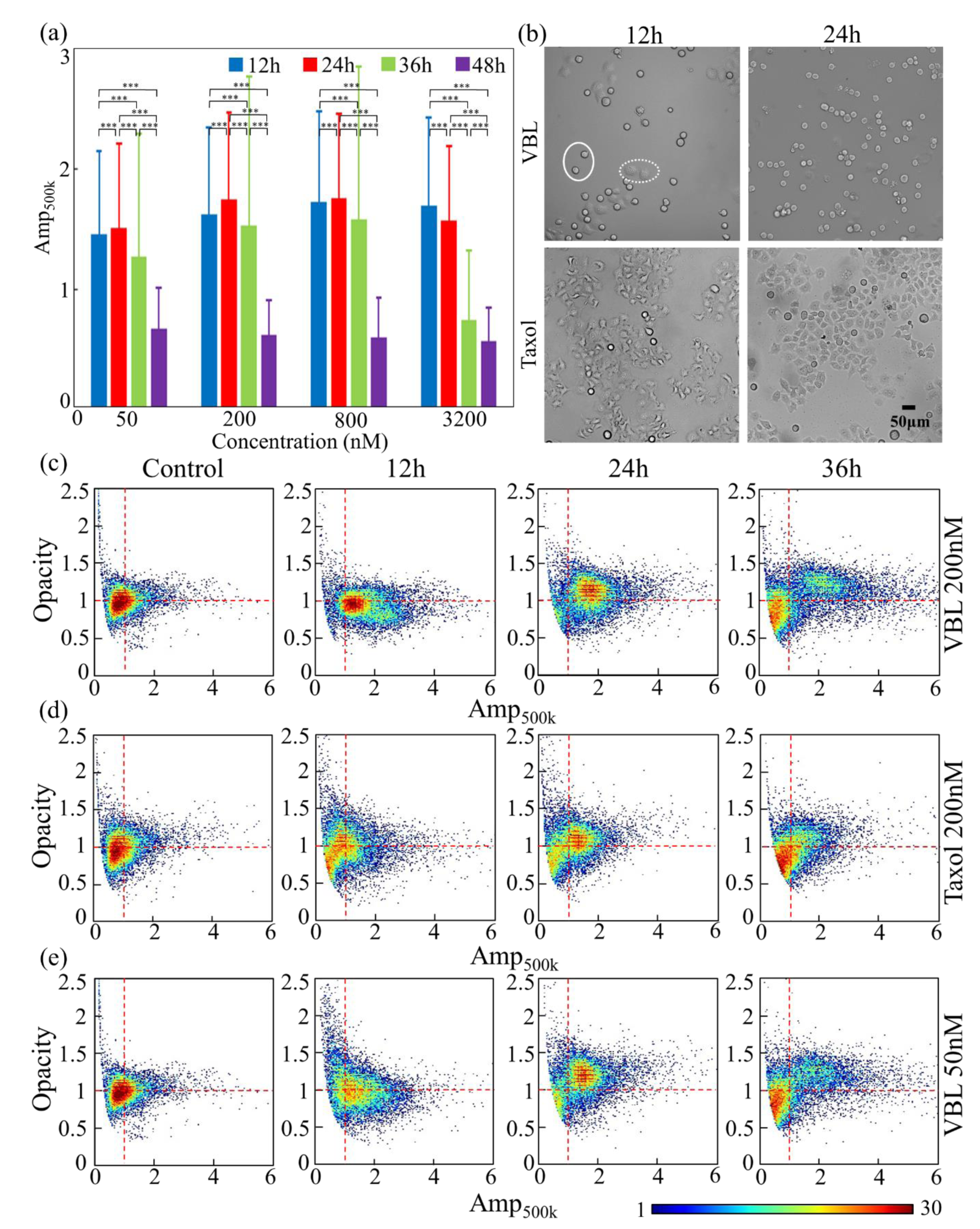
(a) Mean ± s.d. of the ***Amp***_**500*k***_ for VBL-treated cells plotted against the exposure time durations respectively under VBL concentrations of 50-, 200-, 800- and 3200-nM (n = 15,000). (b) Phase contrast images of the VBL- and taxol-treated cells at 50nM concentration and exposure time durations of 12h and 24h. Outlined cells: representative cells exhibiting rounding (solid circle) and prerounding (dashed circle) morphologies. (c-e) Scatter plots of opacity versus ***Amp***_**500*k***_ for the untreated cells as well as the cells post treatment of 12-, 24- and 36-h by (c) 200 nM VBL, (d) 200nM taxol and (e) 50 nM VBL. Each scatter plot contains 15,000 cell events. The color bar designates the cell density at each data point.

Besides the more pronounced volume increase, VBL also induced more progressive apoptosis as indicated by the cell volume shrinkage after 24 h for the low concentrations of 50 nM and 200 nM. Particularly, cells respectively treated by 50 nM/200 nM VBL for 48 h had a mean ***Amp***_**500*k***_ ∼ 0.65 (Fig. 6(a)), indicating that the cell volume is much less than that of the control group (***Amp***_**500*k***_ **≈ 1**). In comparison, cells treated with taxol of the same conditions still had the volume (***Amp***_**500*k***_ ∼ 1.11 in Fig. 3(b)), remaining larger than that of the untreated group. These impedance features were also verified by the images taken for 48 h treated cells respectively at 50- and 200-nM. Cells post VBL exposure (Fig. S6, Column 2 & 3, Row 4) displayed more prominent membrane blebbing as well as fragmented bodies as induced by apoptosis than those treated by taxol (Fig. S5, Column 2 & 3, Row 4).

On the other hand, the cell opacity incorporating the high frequency information also verified a more significant progression of the VBL-treated cells than taxol-treated ones for the low concentration range. The scatter plots of the correlated opacity versus ***Amp***_**500*k***_ in Fig. 6(c-d) compared the dynamics from 12 h to 36 h between VBL-(Fig. 6(c)) and taxol-(Fig. 6(d)) treated Hela cells at 200 nM concentration. Initially at 12 h, the scattered cluster of VBL-treated cells exhibited a more apparent shift to gained ***Amp***_**500*k***_, which corresponded to a larger fraction of cells experiencing volume growth due to the drug-induced mitotic arrest. Such difference can also be easily recognized by comparing the inspection images in Fig. S5 (Column 3, Row 1) and Fig. S6 (Column 3, Row 1). Later at 24 h, both the scatter plots of VBL- and taxol-treated cells displayed resolved subpopulations respectively in the up-right and the down-left regions. Yet, the down-left subpopulation of the VBL-treated cells at 24 h became more enriched than that at 12 h. This was induced by the back-flow of the volume-expanded cells, part of which underwent apoptosis with increased exposure time to 24 h and exhibited decreased opacity and electrical volume. In contrast, the 200 nM taxol-treated cells in the 24 h plot showed a reduced down-left subpopulation than 12 h plot (Fig. 6(d)), with more cells shifted to the other subpopulation in the up-right region because of expanded volume and diluted cytoplasm. The inspection images in Fig. S5-6 (Column 3) agree well with such a trend. Most of the taxol-treated cells at 24 h appear rounding with plump bodies and smooth surfaces, as a sign of G2/M arrest, whereas the majority of the VBL-treated cells started to show irregular and wrinkle appearance of their round-up body and a small portion even exhibited blebbing membranes, indicating that apoptosis occurred.

The back-flow of the taxol-treated cells to the down-left region (Fig. 6(d)) occurred later at 36 h due to apoptosis, whereas the VBL group at the same time point (Fig. 6(c)) already had the majority of the cells events undergoing apoptosis and exhibited even more distinguishable subpopulations. The scatter plot of opacity against ***Amp***_**500*k***_ for the VBL-treated cells at a lower concentration of 50 nM (Fig. 6(e)) exhibited a similar trend with the 200 nM VBL treatment, except that the ***Amp***_**500*k***_shift at 12 h, as well as the subpopulation separation at 24 and 36 h appeared less significant due to the decreased drug concentration.

Statistical analysis of cell populations treated by taxol and VBL at 200nM showed significant difference for both the opacity and ***Amp***_**500*k***_ at each time point (Table S4). Flow cytometry validation of the 200nM VBL-treated cells against increased exposure time has been carried out. The cell cycle and apoptosis plots derived from flow cytometry confirmed the G2/M arrest and increased cell apoptosis post VBL treatment, as shown by a representative group post 24h treatment in Fig. S6(a-b) (the 3^rd^ column). The overall trends against exposure time of cell fractions under G2/M arrest as well as under apoptosis measured by our MIC device also aligned well with that estimated by flow cytometry, as shown in Fig. S6(d).

## Conclusion

An AgPDMS-based MIC device was developed for single-cell analysis of tumor cells that exhibit progressive cell states as a response to anticancer drug treatment at varied concentrations and exposure time duration. Hela cells that have been well-documented to exhibit mitotic arrest at the G2/M phase followed by apoptosis after treatment of anti-tubulin agents, i.e., taxol and vinblastine, were respectively exploited here as models for the impedance cytometry. High throughput analysis of 10,000∼15,000 cells simultaneously at dual frequencies was performed under each treatment condition. The derived results, including the amplitude and phase at low frequency and the high-frequency opacity, respectively revealed increased electrical volume and membrane capacitance as well as reduced cytoplasm conductivity for cells under mitotic arrest as compared to the untreated ones. On the other hand, cells under apoptotic state displayed opposite trends regarding these impedance parameters. Moreover, the apparent electrical volume growth accompanied by declined cytoplasm conductivity indicated here for cells under mitotic arrest offered valuable support, from the aspect of impedance characteristics, for the long-time debating hypothesis of “osmotic swelling”, as a mechanism for mitotic volume increase and the driven force for mitotic cell rounding. Further, the results identified the more potent drug effect of the vinblastine than taxol when both were applied to Hela cells, especially at low concentrations of 50-or 200-nM.

Compared to the existing MIC device, this work implements an innovative way for the integration of AgPDMS microelectrodes with PDMS sidewall blocks, and enabled a sacrificial-layer-free replica molding process being even simpler and lower cost. More importantly, our work gained the benefit of impedance cytometry for assessment of tumor cell states, besides the viability assay as has been previously demonstrated, for cells post chemotherapeutic drug treatment at label-free and high throughput manner.

## Author Contributions

Xiaoxing Xing and Yuan Luo supervised the entire project. Xiaoxing Xing, Yuan Luo and Duli Yu designed the experiments. Xinlong Yang processed the data, conducted part of the experiments and wrote the manuscript. Ziheng Liang and Xueyuan Yuan performed the experiments and processed data. Yao Cai provided expertise in cell culture and handling.

## Conflicts of interest

There are no conflicts to declare.

## Acknowledgements

This work is supported by the National Key R&D Program of China (Grant No. 2021YFF1200300), the Major Research Instrument Program of Chinese Academy of Sciences (Grant No. E24CYA1001), the National Natural Science Foundation of China (Grant No. 61804007), the Fundamental Research Funds for the Central Universities (No. XK1802-4), the Research Funds to the top scientific and technological innovation team from Beijing University of Chemical Technology (Grant No. BUCTYLKJCX06). We thank Miss. Flora YU for the kind proofreading of the manuscript.

## Notes

### Competing Interest Statement

The authors have declared no competing interest.

