## Supplementary materials for "Single-cell impedance cytometry of anticancer drug-treated tumor cells exhibiting mitotic arrest state to apoptosis using low-cost silver-PDMS microelectrodes†"

**Contents**


**Materials and methods** S-3

Reagents S-3

Data processing S-3

Flow cytometry S-4

**Supporting tables** S-5

**Supporting figures** S-6

**References** S-13

**S1. Reagents**

Dulbecco's modified Eagle medium (DMEM), penicillin/ streptomycin (P/S) and trypsin were purchased from Life Technologies, USA. Fetal bovine serum (FBS) was purchased from Biowest, France. The cell culture medium was DMEM containing 10% FBS and 1% P/S. The drug powder of taxol and VBL were obtained from Solarbio, China. The stock solutions of taxol/VBL were prepared by dissolving the drug powder into Dimethyl sulfoxide (DMSO, Sigma-Aldrich, MO, USA) at 3 µM concentration and were maintained at 4 $^{o}$C in dark. The chemicals used for preparing the phosphate buffered saline (PBS, containing 136.89 mM NaCl, 2.68 mM KCl, 1.47 mM $KH_2PO_4$ and 8.1 mM $Na_2HPO_4$; pH 7.4; conductivity 1.6 S/m) were purchased from Sigma-Aldrich, USA.

**S2. Data processing**

Recorded data in csv file format was processed using a custom software program written in python. Differential signal with its amplitude being no less than 1.5 times of the noise amplitude and its transit time between the counter peaks being shorter than 1 ms was identified as a cell event. 10,000~15,000 events were processed for each experiment. Electric current signal being measured simultaneously at 500 kHz and 10 MHz were recorded in csv files at 57.6k Sa/s sampling rate. Each data point was composed of multiple information such as the real and imaginary parts of the complex current signal, as well as the time point for measurement. The data trace for 10 MHz, in format of complex current amplitude versus time, was analyzed as precursor for differential peak detection. The peaks of 500 kHz sharing the same time index with that of the 10 MHz were then automatically obtained. The detection process of a validated differential peak (i.e. cell event) was showcased in Fig. S3. Firstly, a forward search of the data trace versus time was performed with starting point defined by the time point of $P_{max\_n-1}$, which denoted the maximum peak value of the previous validated event. During the forward search, the first identified local maximum value of the 10MHz amplitude (i.e., a data point with amplitude greater than both the amplitudes of the previous and the next data point on the time axis) was stored temporarily as the maximum peak value $P_{max\_n}$ of the proposed event for validation in the current detection process. Secondly, the minimum peak value was searched backward from the $P_{max\_n}$ until the endpoint as defined again by the $P_{max\_n-1}$ of the previous event. During the backward search, the first detected local minimum value that exhibited peak-to-peak amplitude *Amp* and transit time $t_r$ matching the thresholds (i.e., *Amp* above 1.5 times of the noise amplitude and $t_r$ being shorter than 1 ms) when being paired with the given $P_{max\_n}$ was identified as the minimum peak value $P_{min\_n}$ for the differential signal, which was then recorded as a validated cell event. In case no validated minimum peak value was identified by the endpoint, the backward search was exited and the searching for the next local maximum value was initiated. A txt file containing the validated cell events information was output for further processing in data analysis.

To rule out any inter-experiment variations as well as the effect of the entire circuit system,[1-3] the measured amplitude and phase at each frequency for the drug-treated cells were calibrated respectively using the mean amplitude and phase of the control group with untreated cells measured in parallel in each experiment, where the amplitudes of 500 kHz ($\boldsymbol{Amp_{500k}}$) and 10 MHz ($\boldsymbol{Amp_{10M}}$) were derived by

$$\boldsymbol{Amp_{500k} = Amp_{500k\_0}/\overline{Amp_{500k\_ctl}}} \quad (1)$$

$$\boldsymbol{Amp_{10M} = Amp_{10M\_0}/\overline{Amp_{10M\_ctl}}} \quad (2)$$

and the phase shift at 500 kHz ($\boldsymbol{Phase_{500k}}$) and 10 MHz ($\boldsymbol{Phase_{10M}}$) were calculated by

$$\boldsymbol{Phase_{500k}} = \boldsymbol{Phase_{500k\_0}} - \overline{\boldsymbol{Phase_{500k\_ctl}}} \quad (3)$$

$$\boldsymbol{Phase_{10M}} = \boldsymbol{Phase_{10M\_0}} - \overline{\boldsymbol{Phase_{10M\_ctl}}} \quad (4)$$

in which the subscript "0" denoted the originally measured amplitude or phase, and "$\overline{\cdot}$" designated the mean value of the amplitude or phase from the control group. For opacity calibration, the opacity for drug-treated groups and control groups was first calculated by taking the ratio between the calibrated $\boldsymbol{Amp_{10M}}$ and $\boldsymbol{Amp_{500k}}$, and then the opacity was also normalized with the mean opacity value from its corresponding control group.

Low frequency amplitude ($Amp_{500k}$) and phase shift ($Phase_{500k}$) were respectively used for analysis of cell electrical volume and membrane properties (e.g. the membrane capacitance) regarding the varied cell progression states post drug treatment. Opacity given by the amplitude ratio between high and low frequencies ($Amp_{10M}/Amp_{500k}$) was independent of cell volume and position, and thus was used as an indicator of dielectric parameter change of cell interior (e.g. cytoplasm conductivity $\sigma_c$). Scattered plots, histograms as well as mean value with standard deviation were generated for data analysis. Students' t-test was performed to assess the difference between the measured data populations, with the value of $P \leq 0.05$(*) and $P \leq 0.001$(***) designating statistically significant and highly significant, respectively. P value greater than 0.05 denoted not significant (ns).

**S3. Flow cytometry**

Validations of cell volume change as well as the cellular states of G2/M arrest and apoptosis post drug treatment were carried out using flow cytometry (MoFlo XDP, Beckman Coulter, CA, USA). The data was acquired and analyzed by Summit software (Beckman Coulter, CA, USA). Cell sizing and apoptosis were measured upon cell harvesting. The cell size was evaluated by the pulse height of the forward scatter signal (FSC-H). Cell apoptosis was measured by Annexin V-FITC/Propidium Iodide (PI) stain protocol using an Apoptosis Detection Kit (BA00101, Bioss, China). Cell clusters exhibiting "Annexin V-FITC$^{+}$/PI$^{-}$" and "Annexin V-FITC$^{-}$/PI$^{-}$" designated the early and late apoptosis stages, respectively. Cell cycle was assessed based on PI-stained DNA content using a Cell Cycle Analysis Kit (BA00204, Bioss, China), before which the cells were fixed using 70% ice-cold ethanol for 2 hours. Cells at G2/M phase were derived by adapting the Watson's pragmatic algorithm using the flow cytometry software.

**Table S1.** Statistical differences of $Amp_{500k}$ between the control group (Ctl) and the drug-treated groups post 12h exposure (corresponding to Fig.3(b) and Fig. 6(a))

| | 50nM Vs. Ctl | 200nM Vs. Ctl | 800nM Vs. Ctl | 3200nM Vs. Ctl |
|---|---|---|---|---|
| Taxol 12h | * | *** | *** | *** |
| VBL 12h | *** | *** | *** | *** |

**Table S2.** Statistical differences analyzed for cell populations post 3200nM taxol treatment for 12h (control, "flatting" and "rounding" groups in Fig.4(a))

| | Flatting Vs. Ctl | Rounding Vs. Ctl | Flatting Vs. Rounding |
|---|---|---|---|
| $Amp_{500k}$ | *** | *** | *** |
| $Phase_{500k}$ | *** | *** | *** |

**Table S3.** Statistical differences analyzed for cell clusters exhibited in Opacity/$Amp_{500k}$ scatter plot post 3200nM taxol treatment (Fig. 5).

| | 12h | 24h | 36h |
|---|---|---|---|
| Blue Vs. red(for opacity) | * | *** | *** |
| Up-right Vs. down-left (for $Amp_{500k}$) | *** | *** | *** |
| Up-right Vs. down-left (for opacity) | *** | *** | *** |

**Table S4.** Statistical differences analyzed for cell populations post taxol and VBL treatment at 200nM as well as for cell populations post 50nM and 200nM VBL treatment (Fig. 6).

| | 12h | 24h | 36h |
|---|---|---|---|
| Taxol Vs. VBL (for $Amp_{500k}$ at 200nM) | *** | *** | *** |
| Taxol Vs. VBL (for opacity at 200nM) | *** | *** | *** |
| 50nM VBL Vs. 200nM VBL (for $Amp_{500k}$) | *** | *** | *** |
| 50nM VBL Vs. 200nM VBL (for opacity) | * | *** | *** |

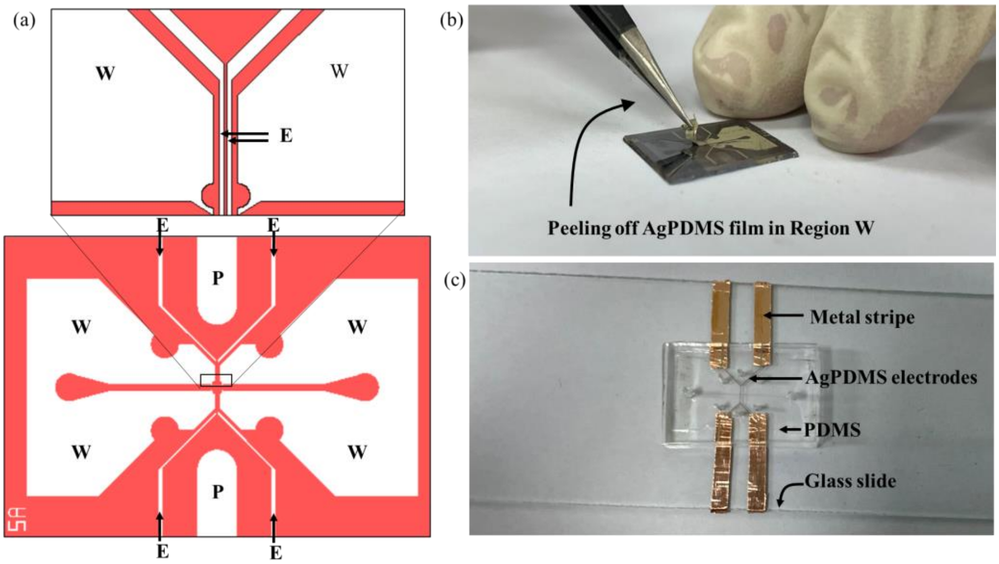


**Fig. S1** (a) Photomask design illustrating the active and passive regions on the device. Region E: microelectrodes (active region). Region W: flow sidewalls (passive region). Region P: supporting posts (passive region). (b) Peeling off of the cured AgPDMS film from the passive region W. (c) A device photo taken after plasma bonding with glass slide and before mounting onto PCB board. The device layer is composed of PDMS passive blocks and AgPDMS electrodes that connect the cooper metal stripes on the glass slide.

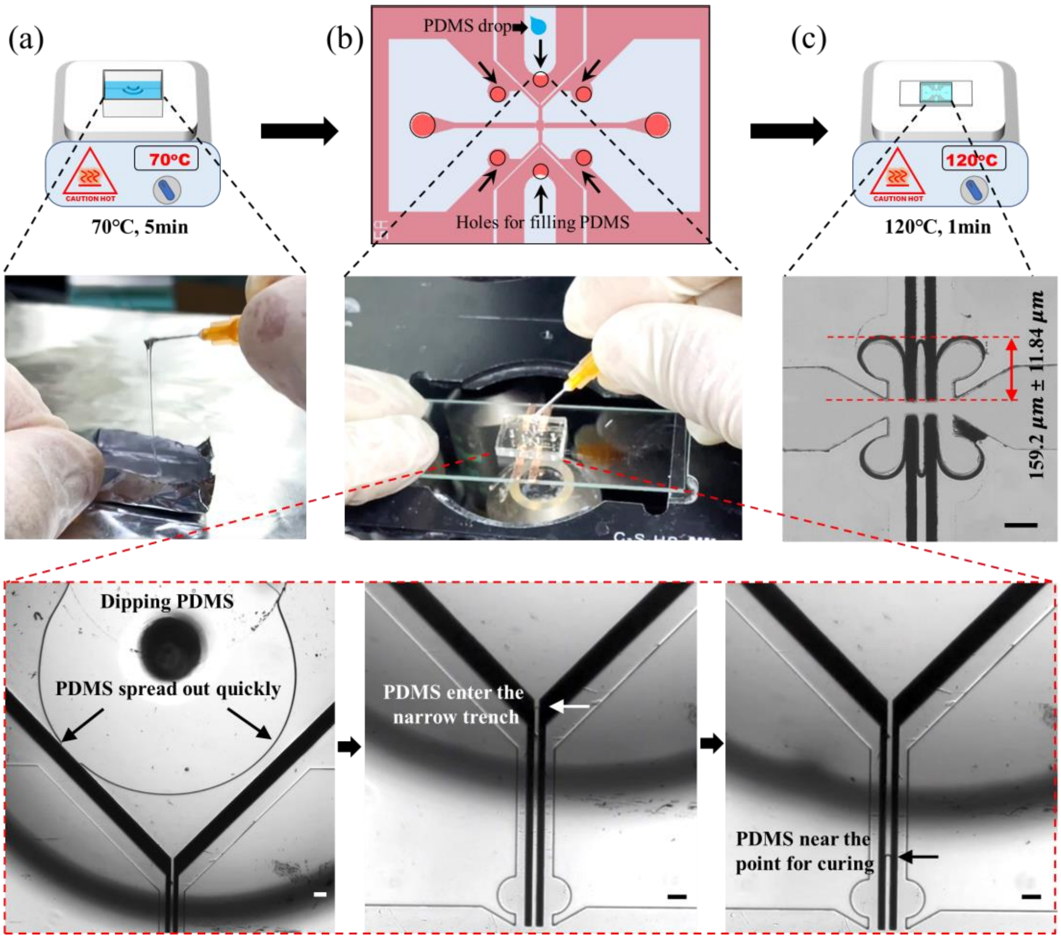


**Fig. S2** Capillary filling of the half-cured PDMS. (a) Pre-heating of the freshly mixed PDMS (70°C, 5min). (b) Dipping of PDMS into the ports by a needle. Bottom panel: sequential micrographs showing the PDMS capillary filling in the middle insulation trench. Scale bar: 80μm. (c) Curing of the filled PDMS (120°C, 1min) and micrograph of a representative device being symmetrically filled. The end point for filled PDMS in the narrower middle trench can be controlled with a length of 159.20μm±11.84μm (n=10) away from the electrode digit ends. Scale bar: 80μm.

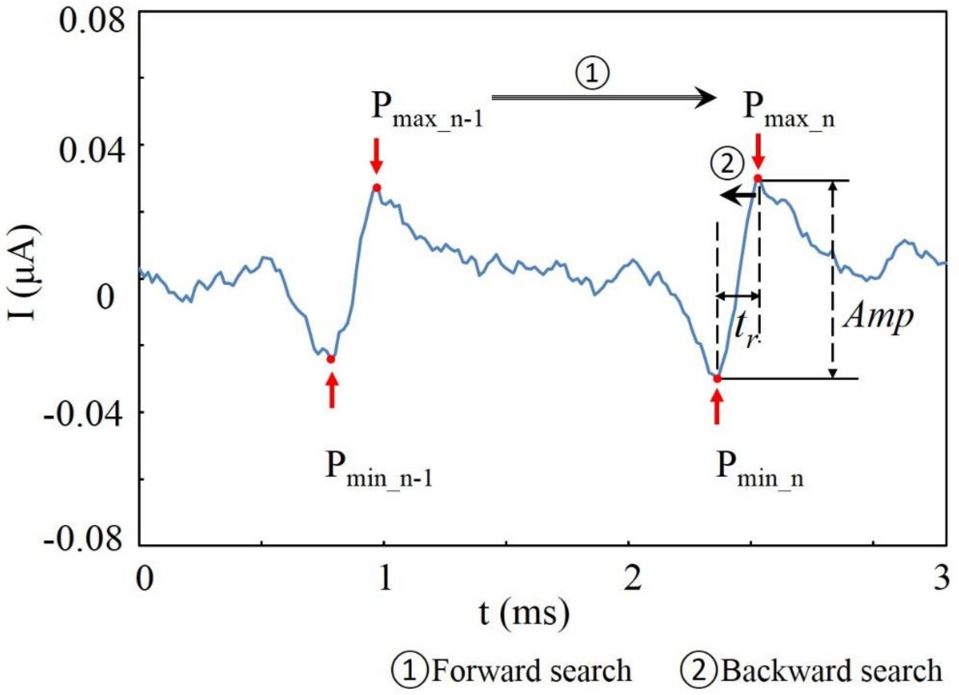


**Fig. S3** 10 MHz amplitudes versus time being partially taken from the recorded data trace illustrating the signal features as well as of the peak detection process for validated cell event. ($P_{min\_n-1}$, $P_{max\_n-1}$) and ($P_{min\_n}$, $P_{max\_n}$) denote the differential peaks of two sequential cell event numbered with n-1 and n in the subscript, and the subscripts max/min refer to the highest/lowest peak of the differential signal. *Amp* and $t_r$ denote the amplitude and the transit time of the differential peaks, respectively.

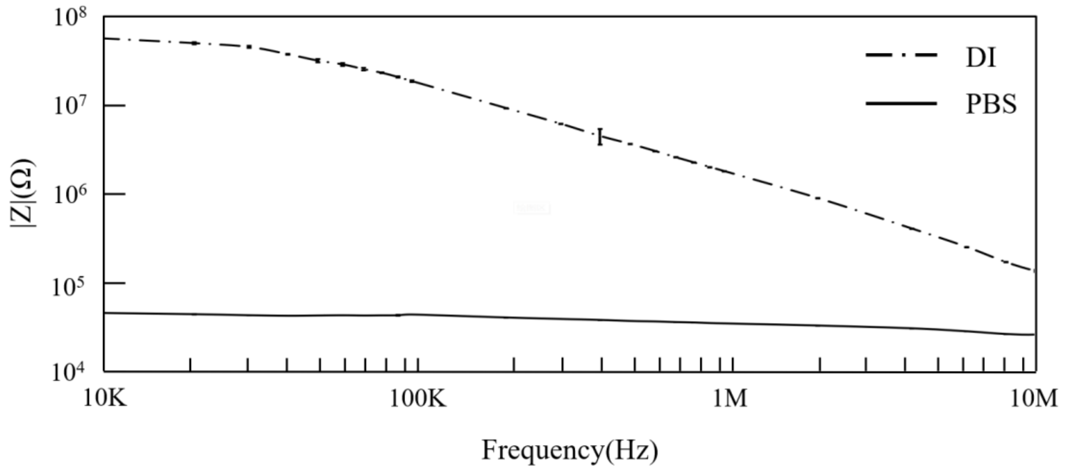


**Fig. S4** Impedance spectra measured using four-probe methods by the impedance spectroscope (HF2IS, Zurich Instruments, Zurich, Switzerland). The measurement was performed between one pair of electrodes with the channel filled with deionized (DI) water and PBS. Data symbols and error bars denote mean ± s.d. (n = 3).

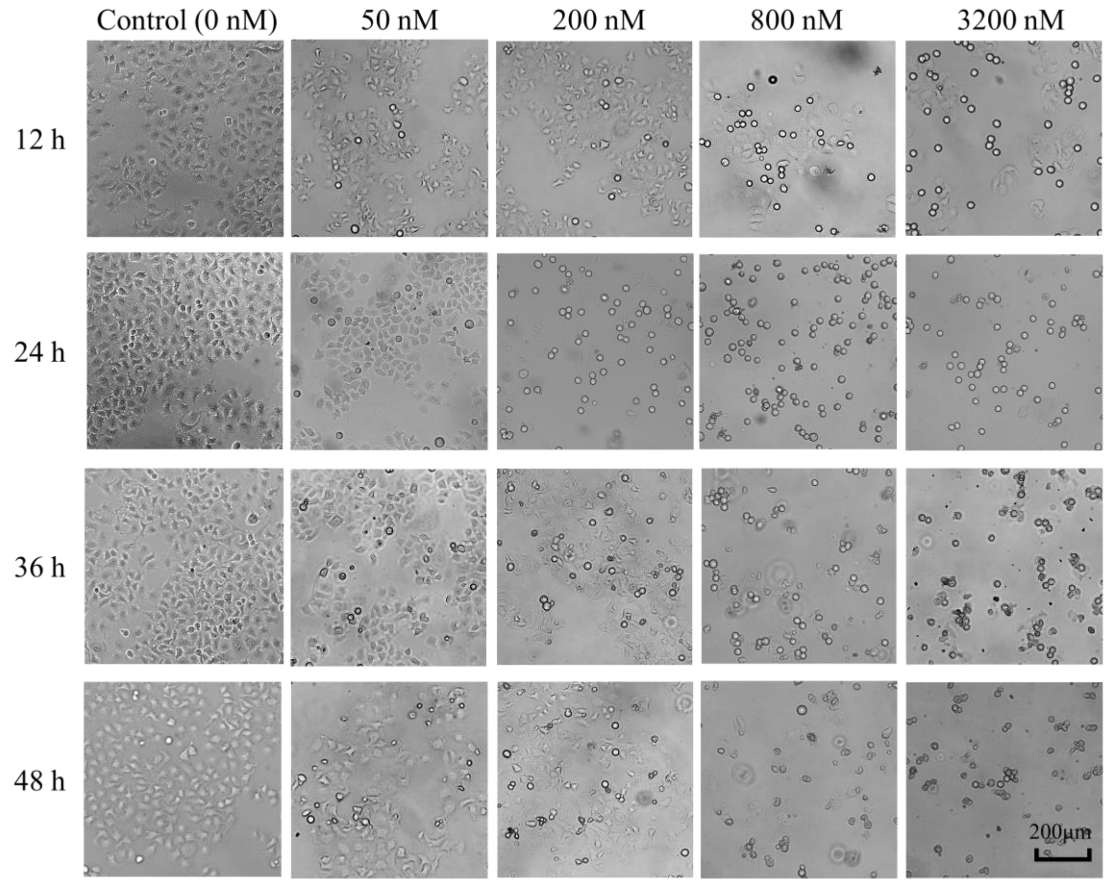


**Fig. S5** Micrograph gallery of Hela cells post taxol treatment respectively at drug concentration of 0 (control), 50 nM, 200 nM, 800 nM and 3200 nM (each row) for different time length of 12 h, 24 h, 36 h and 48 h (each column).

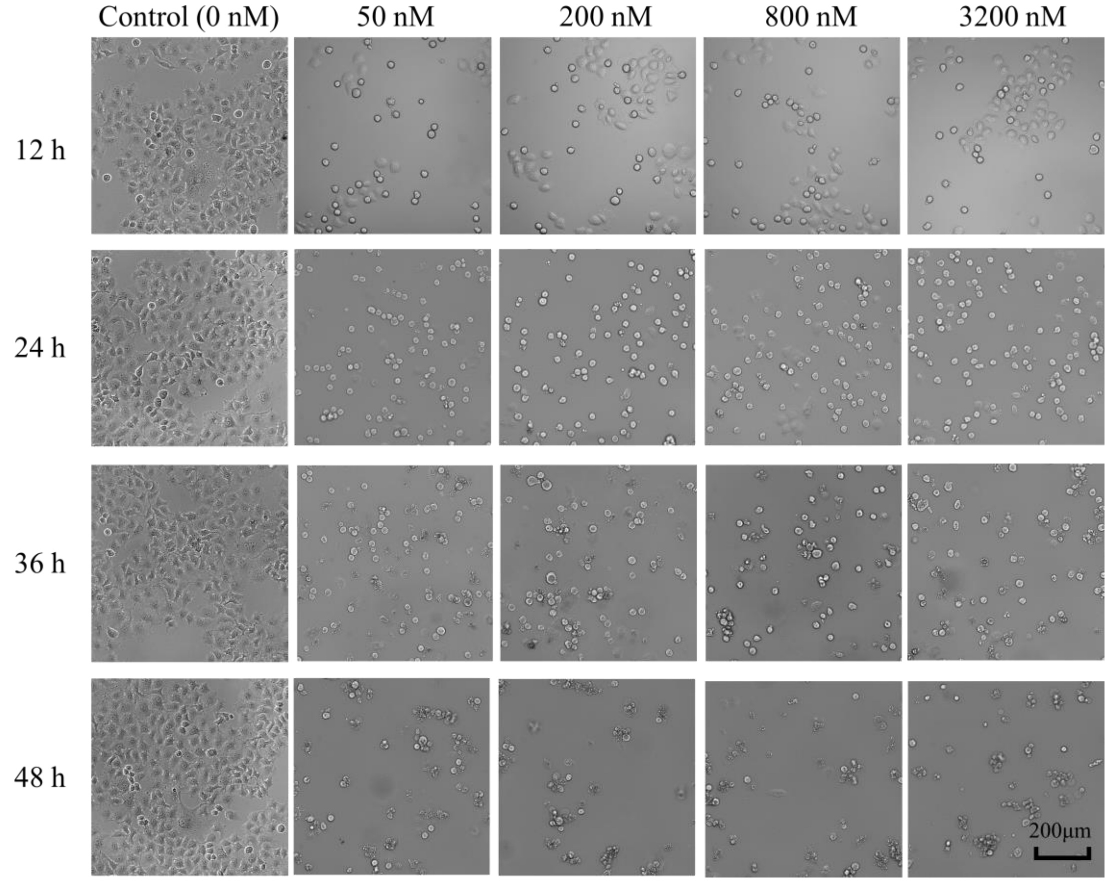


**Fig. S6** Micrograph gallery of Hela cells post VBL treatment respectively at drug concentration of 0 (control), 50 nM, 200 nM, 800 nM and 3200 nM (each row) for different time length of 12 h, 24 h, 36 h and 48 h (each column).

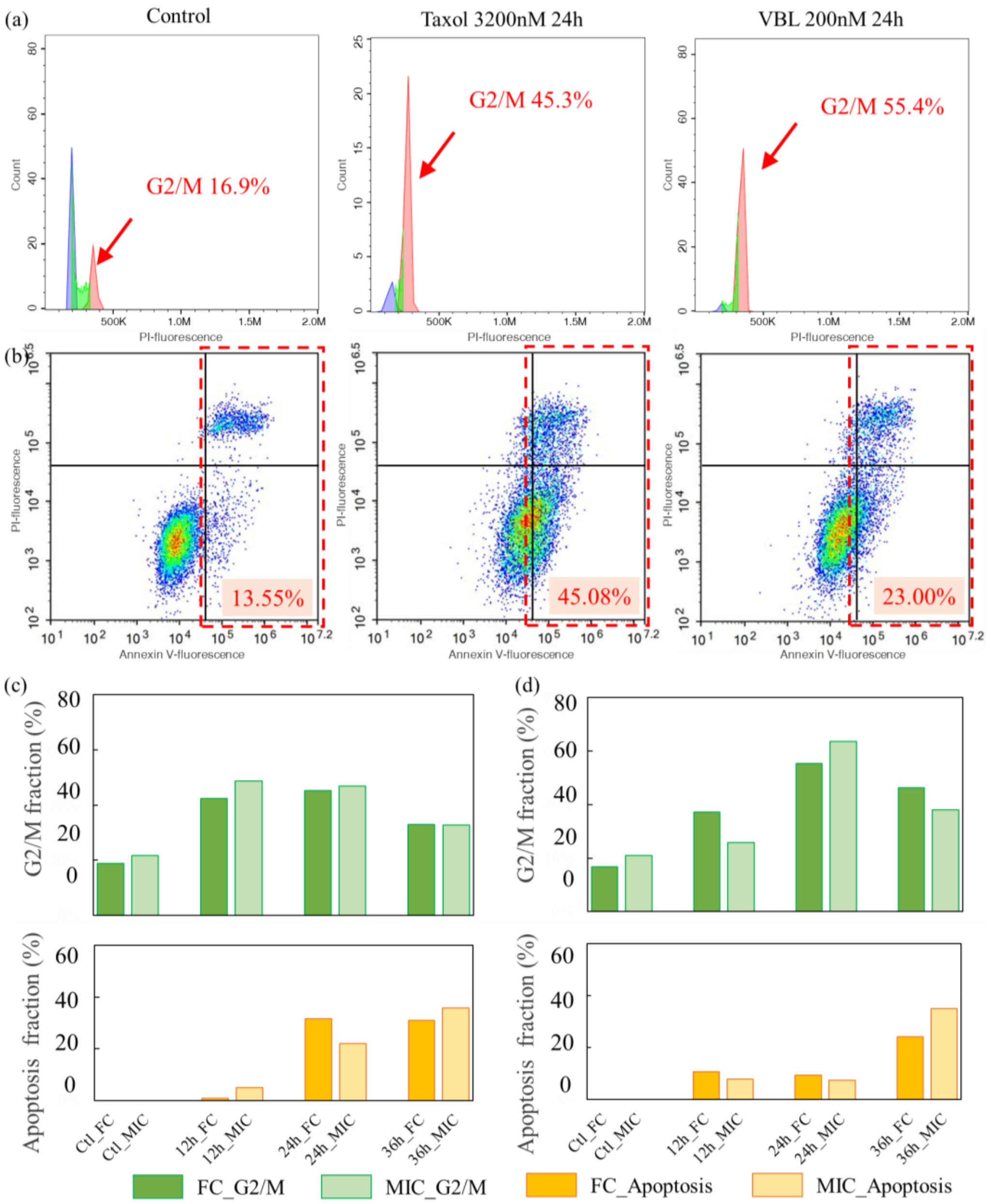


**Fig. S7** (a) PI fluorescence histogram for the control and drug-treated cells. Cell fraction of G2/M phase (pink) are shown. (b) Scatter plot correlating the fluorescence intensity of PI and Annexin V for the control and drug-treated cells. Cell fractions under apoptosis state are outlined by the red dashed rectangles, containing cells in both the early (bottom-right corner) and late (top-right corner) stages. (c-d) Comparison between flow cytometry and the MIC device for the cells fractions under G2/M arrest and apoptosis state they measured, with cells treated by (c) 3200nM taxol and (d) 200nM VBL, respectively.

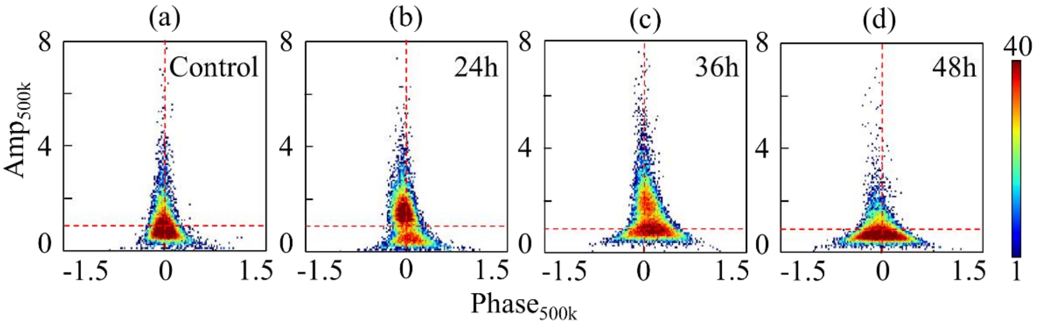


**Fig. S8** Scatter plots of $Amp_{500k}$ versus $Phase_{500k}$ for the (a) control sample with untreated cells and (b-d) the cells treated by 3200 nM taxol respectively at (b) 24 h, (c) 36 h and (d) 48 h. The color bar designates the cell density at each data point.
